# OC_Finder: A deep learning-based software for osteoclast segmentation, counting, and classification

**DOI:** 10.1101/2021.10.25.465786

**Authors:** Xiao Wang, Mizuho Kittaka, Yilin He, Yiwei Zhang, Yasuyoshi Ueki, Daisuke Kihara

**Author notes:** co-first authors (Xiao Wang, Mizuho Kittaka).

## Abstract

Osteoclasts are multinucleated cells that exclusively resorb bone matrix proteins and minerals on the bone surface. They differentiate from monocyte/macrophage-lineage cells in the presence of osteoclastogenic cytokines such as the receptor activator of nuclear factor-κB ligand (RANKL) and are stained positive for tartrate-resistant acid phosphatase (TRAP). In vitro, osteoclast formation assays are commonly used to assess the capacity of osteoclast precursor cells for differentiating into osteoclasts wherein the number of TRAP-positive multinucleated cells are counted as osteoclasts. Osteoclasts are manually identified on cell culture dishes by human eyes, which is a labor-intensive process. Moreover, the manual procedure is not objective and result in lack of reproducibility. To accelerate the process and reduce the workload for counting the number of osteoclasts, we developed OC_Finder, a fully automated system for identifying osteoclasts in microscopic images. OC_Finder consists of segmentation and classification steps. OC_Finder detected osteoclasts differentiated from wild-type and *Sh3bp2^KI/+^* precursor cells at a 99.4% accuracy for segmentation and at a 98.1% accuracy for classification. The number of osteoclasts classified by OC_Finder was at the same accuracy level with manual counting by a human expert. Together, successful development of OC_Finder suggests that deep learning is a useful tool to perform prompt and accurate unbiased classification and detection of specific cell types in microscopic images.

## Introduction

Bone homeostasis is maintained with the balance between bone resorption by osteoclasts and bone formation by osteoblasts, which are tightly coordinated with each other.^1^ Osteoclasts are highly specialized bone-resorbing cells that are differentiated from monocytes/macrophages lineage cells and they play a critical role in various physiological events including bone development, bone repair, and regulation of mineral balance.^1,2^ Excess osteoclast activity will cause bone loss in a variety of pathological conditions such as osteoporosis, rheumatoid arthritis, periodontitis, multiple myeloma, and metastatic cancer. On the other hand, impaired osteoclast activity results in a pathological condition called osteopetrosis characterized by life-threatening bone fragility due to increased bone density.^3,4^ For example, osteoporosis suffers more than 200 million people worldwide, causing a huge socio-economic burden.

Because of the biological importance, osteoclasts have been one of the foci in bone biology. *In vitro* osteoclast differentiation is induced by the stimulation of their progenitor cells with macrophage colony-stimulating factor (M-CSF) and the receptor activator of nuclear factor-κB ligand (RANKL).^1,3^ Differentiated osteoclasts are distinguishable from their progenitor cells by their unique characteristics of multinuclearity and positivity for tartrate-resistant acid phosphatase (TRAP).^3^ Since the establishment of osteoclast culture methods^5^, *in vitro* osteoclast differentiation assays have been extensively used to quantitate and compare the capacity of the progenitor cells for differentiating into osteoclasts. In the assay, the number of TRAP-positive multinucleated cells on culture dishes are manually counted by eyes as osteoclasts by multiple independent examiners. However, the identification of osteoclasts by human eyes does not always secure objectivity and reproducibility. Thus, automated methods for counting osteoclasts have been long awaited.

Here, we developed OC_Finder, a fully automated osteoclast counting system on microscopic images. OC_Finder identifies and segments cells in microscopic images and classifies each cell image into TRAP+ multinucleated osteoclasts and non-osteoclasts. Segmentation is performed with the Otsu’s binarization method^6^ combined with morphological opening and the watershed algorithm^7,8^. The classification of cell images is performed via deep learning, specifically using a convolutional neural network (CNN).

Deep learning has been widely adopted in different biological and medical science areas ^9–13^ for classifying cells in microscope images. However, existing methods have some limitations. For most methods^10–13^, input images need to be manually processed to contain only one cell, or to have cells manually marked to be classified. For other methods ^9^, multi-modal data needs to be prepared as input to help classification. In contrast, in our work we carefully designed the watershed algorithm to segment cell images, which enabled a fully automated framework for cell detection and classification. Unlike existing segmentation methods^14,15^ that need pixel-wise labeling for training, our approach only needs the position of the center of the cells because we perform segmentation in the initial step of the procedure. In CNN, we adopted a teacher-student model ^16^ and image data augmentation techniques for training, which yielded a high accuracy.

There are two recent related works^17,18^ that developed software to detect osteoclasts. The foremost important difference to note is that these two works did not release the datasets they used and their software to the public. Thus, we were unable to compare with their methods and users will not be able to use their methods either. In contrast, the code of our OC_Finder and the dataset we collected are fully released to the public so that biologists can use the software. The dataset and the code will also assist computational biologists to develop new methods. In addition, each of them has notable differences from the current work. The work by Cohen-Karlik et al^17^ used a different neural-network framework to detect cells and classify osteoclasts. Their network outputs bounding boxes of cells while OC_Finder segments the cell region boundaries. We can also see that OC_Finder would be easier to apply to other types of cells, because cell segmentation is performed with an image processing technique that does not need particular training. The second article^18^ provides a tutorial on how to use a commercial software for identifying osteoclasts. Since the software is for general purpose of cell classification, to use the software, users need to prepare a dataset by manual annotation and train a neural network by themselves using the prepared dataset, which may not be an easy task for biologists. The target of the analysis is also different; the pipeline is for cell identification in vivo on histology while OC_Finder is for osteoclast counting in vitro.

OC_Finder achieved 99.4% accuracy in segmentation and 98.1% accuracy for classification. The number of osteoclasts classified by OC_Finder was at the same level as counting by eye. Together, the successful development of OC_Finder suggests that deep learning is a useful tool for performing prompt and accurate identification and classification of cells with characteristic morphological features in microscopic images with no bias. This approach may be applied to classify non-cellular objects. OC_Finder is available at http://github.com/kiharalab/OC_Finder. The dataset used in this work is also made freely available at https://doi.org/10.5281/zenodo.5022015.

## Results

### Cell image dataset collection

Osteoclasts exhibit a variety of morphologies depending on the conditions. In order to train a neural network that can recognize a wide range of morphologies of osteoclasts, it is important to include images of osteoclasts from many distinct conditions in the training dataset. We obtained osteoclast images from cultures with several different cytokine stimuli, including different concentrations of RANKL (25 and 50 ng/ml), combinations of RANKL (50 ng/ml) with IL-1β (10 ng/ml) or TNF-α (100 ng/ml), and with osteoclast precursors from mice of different genotypes, *Sh3bp2^+/+^* and *Sh3bp2^KI/+^*. The osteoclast precursors from *Sh3bp2^KI/+^* mice form more and bigger osteoclasts.^19^ IL-1β and TNF-α are pro-inflammatory cytokines that are known to support osteoclast differentiation.^20^ With these variations of conditions, we were able to obtain images of a wide variety of osteoclast morphologies (Fig. 1a).

**Fig. 1.**
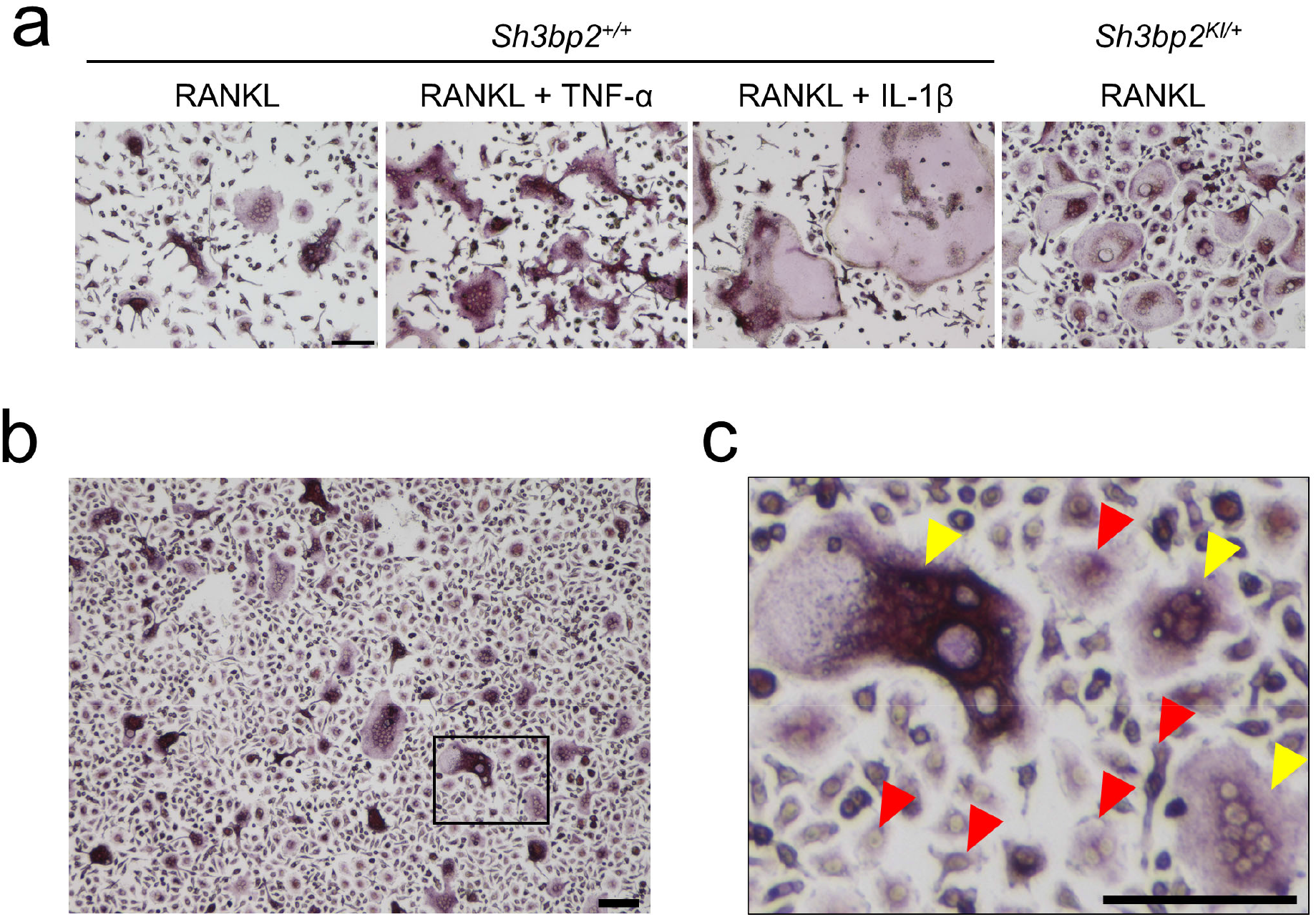
Dataset of cell images of osteoclasts. **a**, Examples of images of various forms of osteoclasts obtained under different conditions. The concentrations of each cytokine in the culture media were as following; RANKL: 50 ng/ml, TNF-α: 100 ng/ml, IL-1β: 10 ng/ml. **b**, an example of captured microscopic images of osteoclast culture. **C**, a magnified image of the boxed area in panel b, showing the examples of induced osteoclasts and non-osteoclasts. The cells which are positive for TRAP-staining and have more than 3 nuclei were identified as osteoclasts (yellow arrowheads), while all other cells which do not satisfy the criteria were regarded as non-osteoclasts (red arrowheads). Black bar = 100 μm.

We obtained 458 microscopic images of osteoclast culture in total. In each image, we manually located the osteoclasts and non-osteoclasts in the images (Fig. 1b and 1c). We generated two datasets from these images. The first dataset was for testing the segmentation accuracy of OC_Finder (the segmentation dataset). The second dataset was to examine the classification accuracy of the method (the classification dataset).

For the segmentation dataset, we selected 10 microscopic images of different culture conditions. The culture conditions were as follows: Osteoclast precursors from males or females with the genotype of *Sh3bp2^+/+^* or *Sh3bp2^KI/+^* stimulated with 25 or 50 ng/ml of RANKL; and osteoclast precursors from male wildtype mice stimulated with the combination of 50 ng/ml of RANKL with IL-1β (10 ng/ml) or TNF-α (100 ng/ml). In each of the 10 images, we manually counted all the cells. The number of manually identified cells in an image ranged from 445 to 1823 with a total of 10221.

For the classification dataset, from each of the 458 images we manually identified about 60 cells, only a fraction of cells in an image, so that we could cover a large number of different microscopic images. In total, we located and labeled 13,822 osteoclasts and 13,833 non-osteoclasts. A cell was considered an osteoclast if it is positive for TRAP staining (pink to purple color in Fig. 1) and has more than 3 nuclei and were considered non-osteoclasts otherwise. Among the 458 microscopic images, 373 images (81.4%) were used for training and validation while the rest (85 images) were used for testing OC_Finder. The 373 images were further split into 298 images (79.9%) for training, which included 9,276 osteoclasts and 9,278 non-osteoclasts, respectively, and 75 images (20.1%) for validation, which included 2,219 osteoclast and 2,226 non-osteoclasts, respectively. The 85 testing images included 2,327 osteoclasts and 2,329 non-osteoclasts, respectively.

For evaluating the cell classification task, we manually segmented out cell images of a size of 50×50 pixels centering at the manually assigned center locations from the microscope images. Since the training and the testing sets were taken from different microscope images, there was no overlap between the two sets. When a cell image was used for training, augmentation via one of 12 types of image transformation listed in Supplementary Table 1, such as translation, rotation, changing contrast, was applied with a random magnitude. The augmentation process allows a significantly higher amount of trainable data to be derived from the fixed amount of images present in the training dataset. The details of augmentation are addressed in the Methods section.

### Overall Architecture of OC_Finder

OC_Finder processes a given microscopic image with two major steps: segmentation and classification (Fig. 2a). First, the program identifies cells in the microscopic image and segments them with the watershed algorithm. Next, small cells are removed since they are unlikely to be osteoclasts. Then, the region of each cell is trimmed into the same square image and three colors in the image, RGB, are normalized considering the mean and the variance of the image. Then, the trained deep learning model is applied to all the trimmed cell images and assign labels. In Fig. 2, non-osteoclasts are assigned 0 and osteoclasts are assigned 1. Finally, OC_Finder visualizes the results with labels assigned to all the segmented cells on the original microscopic image.

**Fig. 2.**
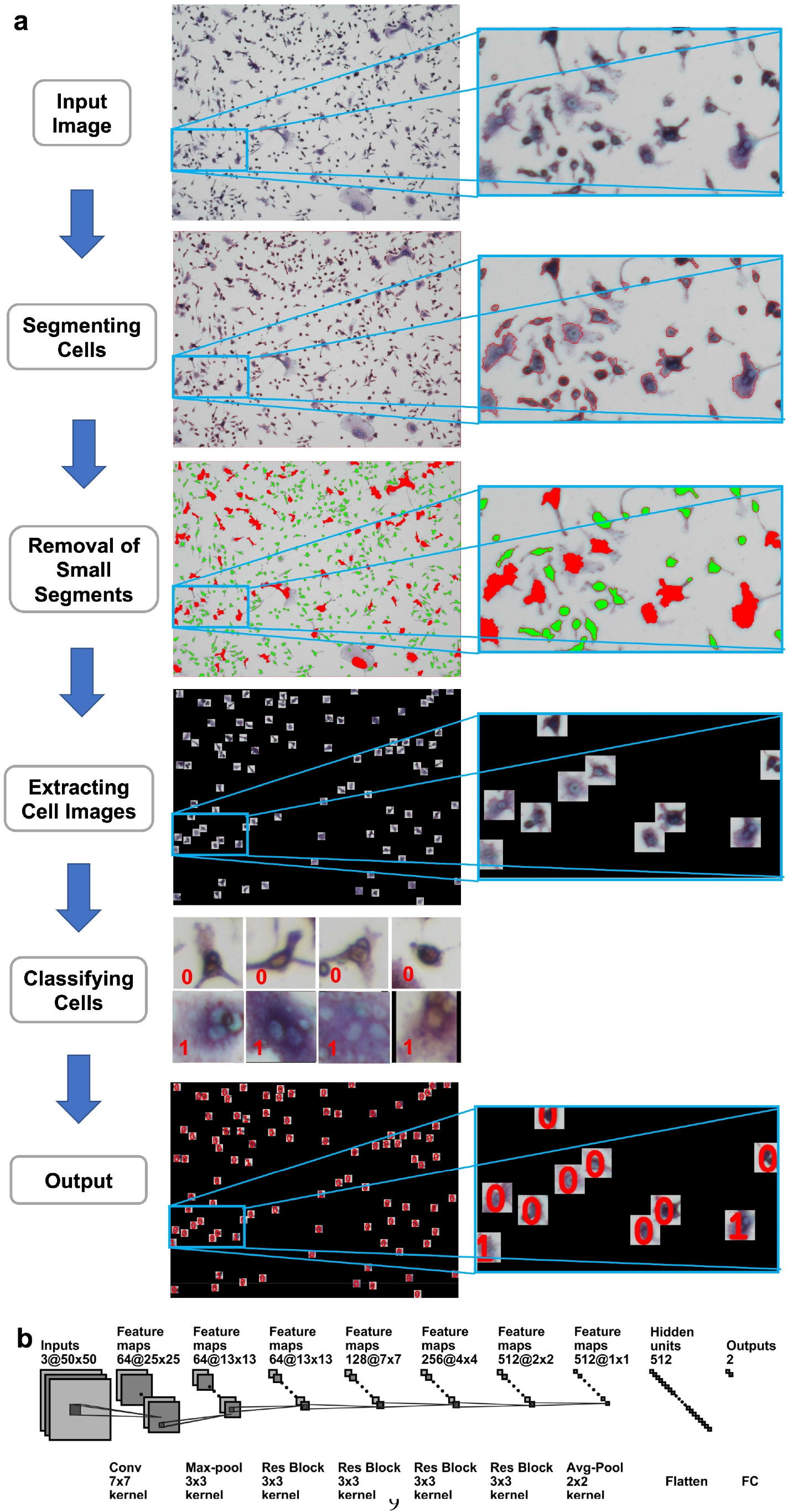
The diagram of OC_Finder. **a.** The workflow of OC_Finder. After cells are segmented, small cells with less than 500 pixels are removed (green) from further downstream analysis because such cells are never osteoclast. Remaining cell images are trimmed to the size of 50×50 pixels, which are input for the deep learning model that classifies it to either non-osteoclast or osteoclast. Finally, OC_Finder will present the microscopic image with predicted labels assigned to identified cells. **b**. The deep learning network architecture for cell classification. The architecture is the same as ResNet-18^21^. Res Block, the residual block, which combines convolution layers, batch normalization, and residual connection (Supplementary Fig. 1). The notation of the layers, for example, 64@25×25 indicates 64 feature maps of 25×25 size and Conv 7×7 kernel represents the convolutional operation with a kernel size of 7×7. Finally, the network outputs the probabilities that the input cell is non-osteoclast or osteoclast.

#### Segmentation

An input microscopic image includes many cells. To start the process, we need to identify and segment cells before applying the deep learning network for cell classification. A microscope image has color values in RGB, which are converted into grayscale values by averaging the three values of Red, Green and Blue. The resultant grayscale microscope images thus depict cellular regions in darker gray than the background. To detect cells, the watershed algorithm is applied, which starts from significantly high dark points (which we term markers) and lower the watershed line to capture neighboring pixels that are included in cells. However, since a naïve application of the watershed algorithm resulted in oversegmentation of cell regions to smaller pieces, we designed the following procedure to reduce markers: First, we applied Otsu’s binarization method^6^ to an input microscopic image, which automatically finds a proper threshold to separate background and the foreground that includes cells to roughly estimate potential cell regions. Next, we removed small or irregular foreground regions by applying a morphological opening operation^22^ with a filter of a 3×3 pixel size, which scans the foreground and removes pixels in the filter from the foreground if the filter is not entirely filled with foreground pixels. This opening operation greatly reduces noise from the foreground. Similarly, we also applied a morphological closing operation^22^ with a 3×3 filter, which scans the input image and converts background pixels in the filter to the foreground if the filter is not entirely filled with background pixels. This process removes small holes and modifies irregular boundary regions in the foreground. In the segmentation process, very large segmented areas can be problematic because they may consist of two or more cells, which cause under-segmentation. To address this problem, we identified potential cell centers in the foreground, which are pixels that are distant from the background. More precisely, a microscopic image, which is now binary labels, foreground and background at each pixel, is re-labeled with the distance to the closest background pixel. Then, pixels with larger distance values than a threshold are selected as markers for the watershed algorithm. Finally, we applied the watershed algorithm using the markers as starting points to segment cells in the input microscopy image. Lowering watershed was stopped once it reached a background pixel or two watershed levels from neighboring cells met.

After segmentation, we removed small segmented regions with less than 500 pixels because they are either a part of a large cell or non-osteoclast for 100% of the cases and would not affect the results of detecting osteoclasts. Then, each cell region is extracted by a square of 50×50 pixels that are placed at the center square of the cell determined by segmentation (Supplementary Fig. 2c). These square images are inputs for the cell classification by deep learning.

### Network Architecture

Fig. 2b shows the neural network architecture of the cell classification model. We used the ResNet-18 architecture^21^. An input is a color cell image in RGB with a size of 50×50 pixels. 64 convolutional filters of 7×7 pixels scan the input with a stride of 2 pixels to capture the local texture pattern of the image. This step results in 64 feature maps of a 25×25 size. Subsequently, a max-pooling layer, four residual block layers (Supplementary Fig. 1) with 64, 128, 256, and 512 residual blocks, respectively, are applied. Then, the output from the last residual block is processed through an average pooling layer to obtain a feature vector. Finally, the feature vector is flattened and passed to a Fully Connected (FC) layer with 512 neurons and activated by a softmax activation function to produce the probability values that the input cell is non-osteoclast or osteoclast.

The network was trained and validated on manually labeled cell images of a 50×50 pixel size in the training set, which contains 11,495 and 11,504 non-osteoclast and osteoclast images respectively, in the 81.4% (357 microscopy images) of the entire microscopic image dataset. During training, cell images were augmented by one of the 12 randomly selected augmentations to increase the number of training views of the same instance. Training was performed with a teacher-student model^16^ where weights of a teacher model were updated with an average of the weights from a sequence of student models of previous iterations while weights of the student model were updated at each iteration.

Due to this weight optimization mechanism, in general, the teacher model is more stable and has higher generalizability and effectiveness in supervised training.

### Segmentation results

First, we discuss the accuracy of the segmentation step. This step corresponds to “Segmenting Cells” in Fig. 2a. Examples of segmentation are shown in Fig. 2a, Fig. 3, and Supplementary Fig. 2. The first two panels in Fig. 2a and Supplementary Fig. 2a and b show an example of an input image and a segmentation result. In the second panel of Fig. 2a and Supplementary Fig. 2b, boundaries of segmentations are shown in red, which correspond well to cells in the image. The detailed segmentation results on the 10 microscopic images in the segmentation dataset are provided in Supplementary Table 2. On average, OC_Finder showed a high detection rate of 99.4% of manually detected cells. There were 80 cells that were missed by OC_Finder in the 10 images. Among them, there was only 1 osteoclast included. On the other hand, OC_Finder detected 3 regions that were not included in the manually detected cells. These three regions were not cells but debris, and they are all removed in the subsequent step of the removal of small regions of less than 500 pixels. The removed debris are shown in Fig. 3a (yellow arrowheads).

**Figure 3.**
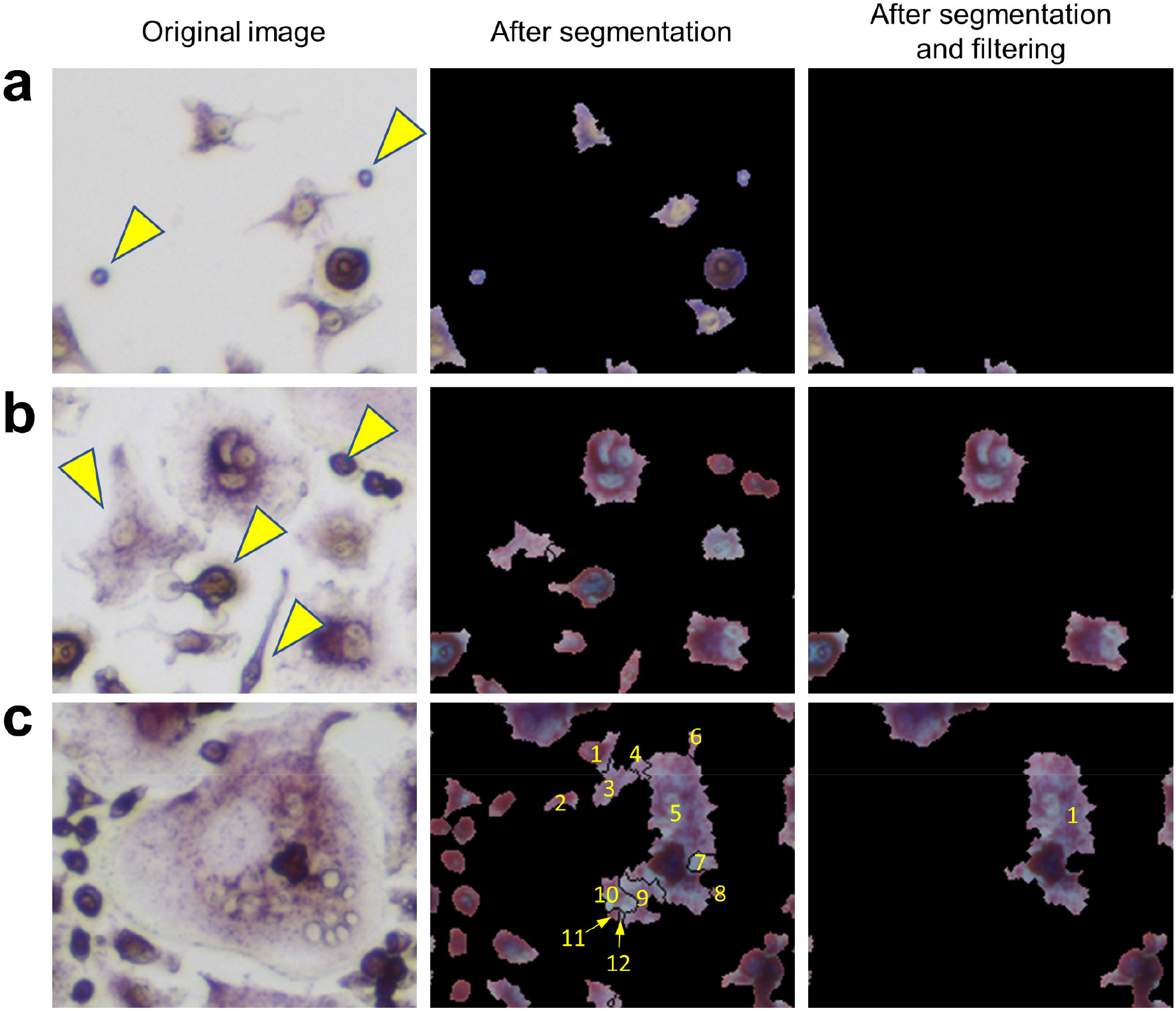
Examples of segmentation and filtering of small regions. Three examples are shown. Left: original images. Middle: After segmentation. Right, after filtering. regions that were filtered out are pointed by yellow arrowheads. **a**, examples of removed small debris. **b,** an example that small regions that correspond to non-osteoclasts were removed by filtering. **c**, an example where a large cell was segmented into multiple regions. A cell was segmented to 12 areas. Out of them, 11 small regions were removed, leaving only one area remaining. This area was later sufficient to correctly classify this cell as an osteoclast.

The step of removing small regions is illustrated in the third panel in Fig. 2a. In the panel, removed segmentations are shown in green, while the remaining large regions are colored in red. Fig. 3b and 3c are additional examples that illustrate the removal of small regions by applying the 500-pixel cutoff. Most of the segmented regions that were removed were non-osteoclasts. Fig. 3b shows a part of a microscopic image that includes removed cells (indicated with yellow arrowheads). Fig. 3c is an example of a different case, where the removal of small regions helped avoid an overlapped counting of large osteoclasts. In this example, a large osteoclast was segmented into 12 pieces but 11 of them were removed by the filtering, leaving only the largest region, which ultimately allowed it to be correctly classified as an osteoclast.

### Cell classification results

Next, we discuss cell classification accuracy. The classification performance was evaluated on the test set of the classification dataset, which includes 2,327 osteoclasts and 2,329 non-osteoclasts, respectively, in 85 microscopic images. Results are summarized in Table 1.

**Table 1.** Cell Classification Accuracy.

**A.** All classification test set.
| Labels\Prediction | Pred. as Osteoclast | Pred. as non-Osteoclast | Total |
| --- | --- | --- | --- |
| Osteoclast | 2274 | 53 | 2,327 (100%) |
| Non-osteoclast | 35 | 2294 | 2,329 (100%) |
| Total | 2309 (98.5%) | 2347 (97.7%) | 4,656 |

**B.** After segmentation was applied to the classification test set.
| Segmented\Prediction | Pred. as Osteoclast | Pred. as non-Osteoclast | Total |
| --- | --- | --- | --- |
| Osteoclast | 2,104 (96.6/90.4%)<sup>a)</sup> | 75 | 2,179 (100/93.6%)<sup>a)</sup> |
| Non-osteoclast | 61 | 2,247 (97.4/96.5%) | 2,308 (100/99.1%) |
| Total | 2,165 | 2,322 | 4,487 (100/96.4%) |

- a) The percentage was computed with two references (denominators). The first percentage was relative to the number of cells that were segmented by the segmentation procedure among all the cells in the classification data set. Thus, 2,179 and 2,308 were used for osteoclast and non-osteoclasts, respectively. The second percentage was computed relative to all the cells in the classification dataset. The percentage values computed in this way were smaller than the former as shown, because 3.6% (4,654-4,487) of cells were not correctly segmented and identified by the segmentation procedure.

A high classification accuracy, 98.1% (2274 + 2294/4,656), was achieved on all the cells in the classification dataset (Table 1A). 97.7% (2274+53/2,327) of osteoclasts and 98.5% (2294+35/2,329) of non-osteoclasts were correctly classified. These results were obtained by the teacher model in the Teacher-Student network we used. In Supplementary Table 3, we compared the current model with other models that used different parameter values (for a smoothing coefficient, α. See Methods). Particularly, the table shows that the current teacher model performed better than the model that did not use the teacher-student architecture.

Using the classification dataset, we have also evaluated the entire pipeline of OC_Finder, where the segmentation and the classification steps were applied sequentially (Table 1B). On this dataset, 96.4% of the cells were segmented correctly. These cells include 93.6% of osteoclasts and 99.1% of non-osteoclasts. 4,351 (2,104 + 2,247) cells were correctly classified. The classification accuracy was 97.0% relative to 4,487 correctly segmented cells. If considering all the 4,656 cells in the classification dataset including miss-segmented cells, the accuracy would slightly drop to 93.4%.

Fig.4 shows examples from the validation process for cell classification by OC_Finder. Fig. 4a shows manually assigned labels to a microscopic image in the classification dataset. Cells marked with red and blue are osteoclasts and non-osteoclasts, respectively. In the classification dataset, only a part of cells are manually labeled as mentioned earlier. In Fig. 4b, classification results by OC_Finder for this microscopic image is shown. As discussed earlier, small regions were not processed as they are not osteoclasts. The remaining panels contain examples of cells classified by OC_Finder. Fig. 4c shows examples of osteoclasts (four images on the left) and non-osteoclasts (right) that were correctly classified by OC_Finder. One can see that identified osteoclasts were stained by TRAP staining and have more than 3 nuclei while non-osteoclasts do not have the properties. Fig. 4d shows the opposite, where OC_Finder misclassified cells. On the left shown are four osteoclasts that were wrongly classified as non-osteoclasts. Nuclei in these cells seem to have unclear boundaries, which might be a reason for the misclassification. The images on the right are non-osteoclasts, which were misclassified as osteoclasts. These four images contain overlapped or adjoining cells that resemble multiple nuclei, which may have confused OC-Finder. Fig. 4e is an interesting case where OC_Finder performed better than the human examiner. This cell has three nuclei but the human examiner thought there were only two and thus classified as non-osteoclast since two nuclei are very close to each other and the boundary is not clear. Despite this difficulty, OC_Finder was able to classify it as osteoclast. The last panel (Fig. 4f) are examples where OC_Finder correctly identified osteoclasts from manually unlabeled cells in a microscopic image.

**Fig. 4.**
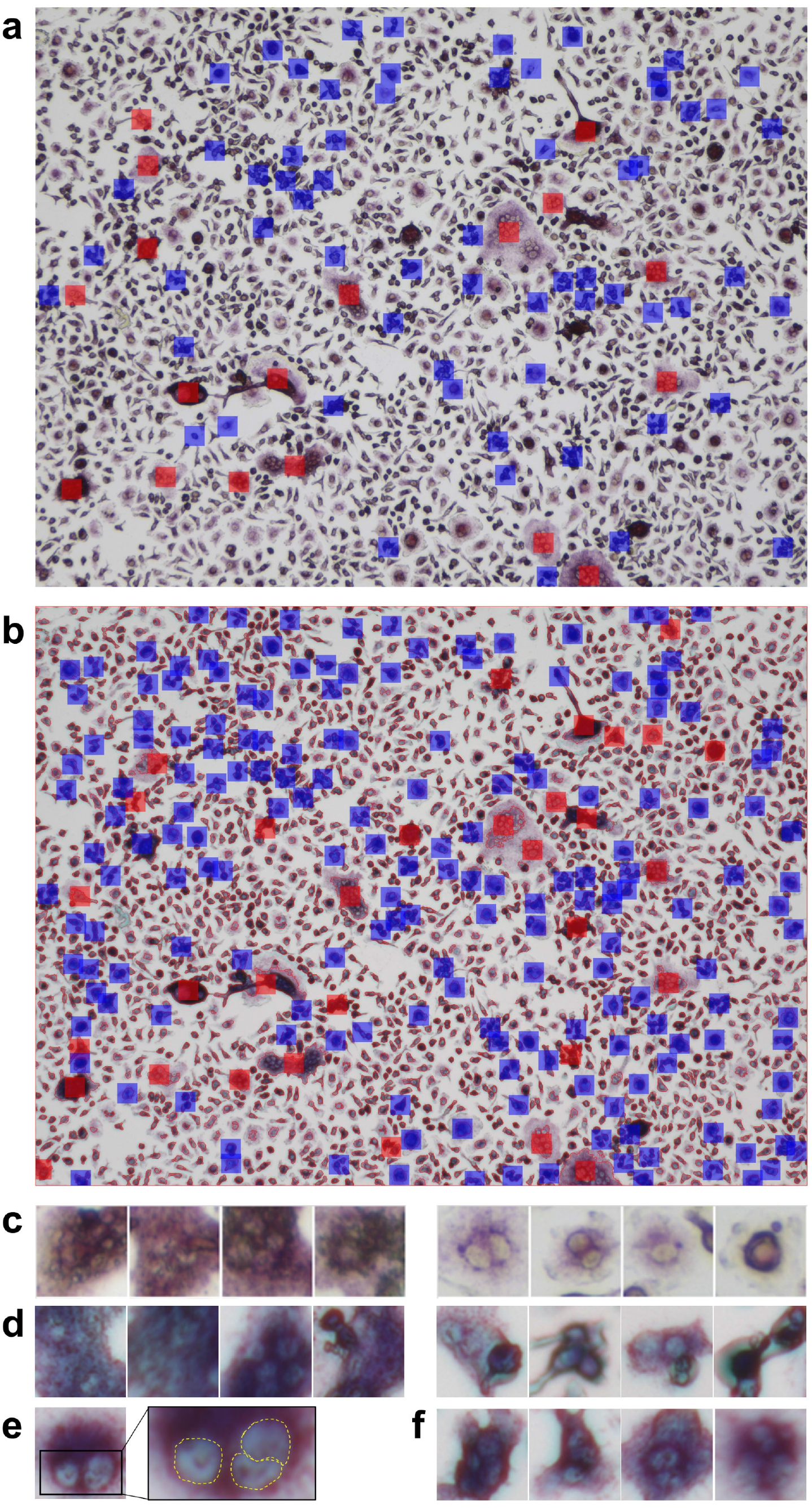
Example of segmentation and classification by OC_Finder. **a.** manually labeled osteoclast and non-osteoclasts in a microscopic image. Red indicates osteoclast and blue indicates non-osteoclast. **b.** osteoclasts and non-osteoclasts detected by OC_Finder for the same image. A red box indicates osteoclast and a blue box indicates non-osteoclast. Cells that are segmented are surrounded by a thin red line. Cells are labeled only when they have a size of 500 pixels or larger. **c.** Examples of osteoclasts (left) and non-osteoclasts (right) images correctly identified by OC_Finder. **d**. The examples of osteoclasts (left) and non-osteoclasts (right) images that were misclassified by OC_Finder. **e**. osteoclast image that was misclassified by manual annotation but correctly classified by OC_Finder. Right panel is the magnified image of the boxed area in the left panel. **f**. Examples of osteoclasts that were not picked by the human examiner during the classification dataset construction and identified as osteoclasts by OC_Finder.

Through this validation process, we confirmed that OC_Finder has a high classification accuracy. We also reaffirmed that humans are prone to error and may occasionally misclassify cell images, in which case OC_Finder can serve as a counteractive measure to human mistakes.

### The performance of the system in a practical situation

Finally, we validated the OC_Finder’s performance in a real-case scenario. Specifically, we examined if OC_Finder could detect an increased osteoclast formation caused by the gain-of-function mutation of *Sh3bp2* (*Sh3bp2^KI/+^*). The gain-of-function of SH3BP2 is known to increase osteoclast formation^19,23,24^. In this experiment, we included cell sources of different gender (male and female), which is an important factor in fields of biology including bone biology,^25^ as well as two concentrations of RANKL (25 and 50 ng/ml) to test whether OC_Finder shows good performance under different experimental designs. Manual counting showed a higher number of osteoclasts in *Sh3bp2^KI/+^* culture (Fig. 5a, the lower panel) than wildtype control without gain-of-function of SH3BP2 (*Sh3bp2^+/+^*) (Fig. 5a, the upper panel) in the four conditions tested (Fig. 5b upper panel, in which result of *Sh3bp2^+/+^* and *Sh3bp2^KI/+^* are presented in white bars and grey bars respectively). These results are consistent with previous reports^19^. The automated counting by OC_Finder also detected the difference between *Sh3bp2^+/+^* and *Sh3bp2^KI/+^* in all the conditions tested (Fig. 5b lower panel) with the same significant *p*-value to the manual counting results.

**Figure 5.**
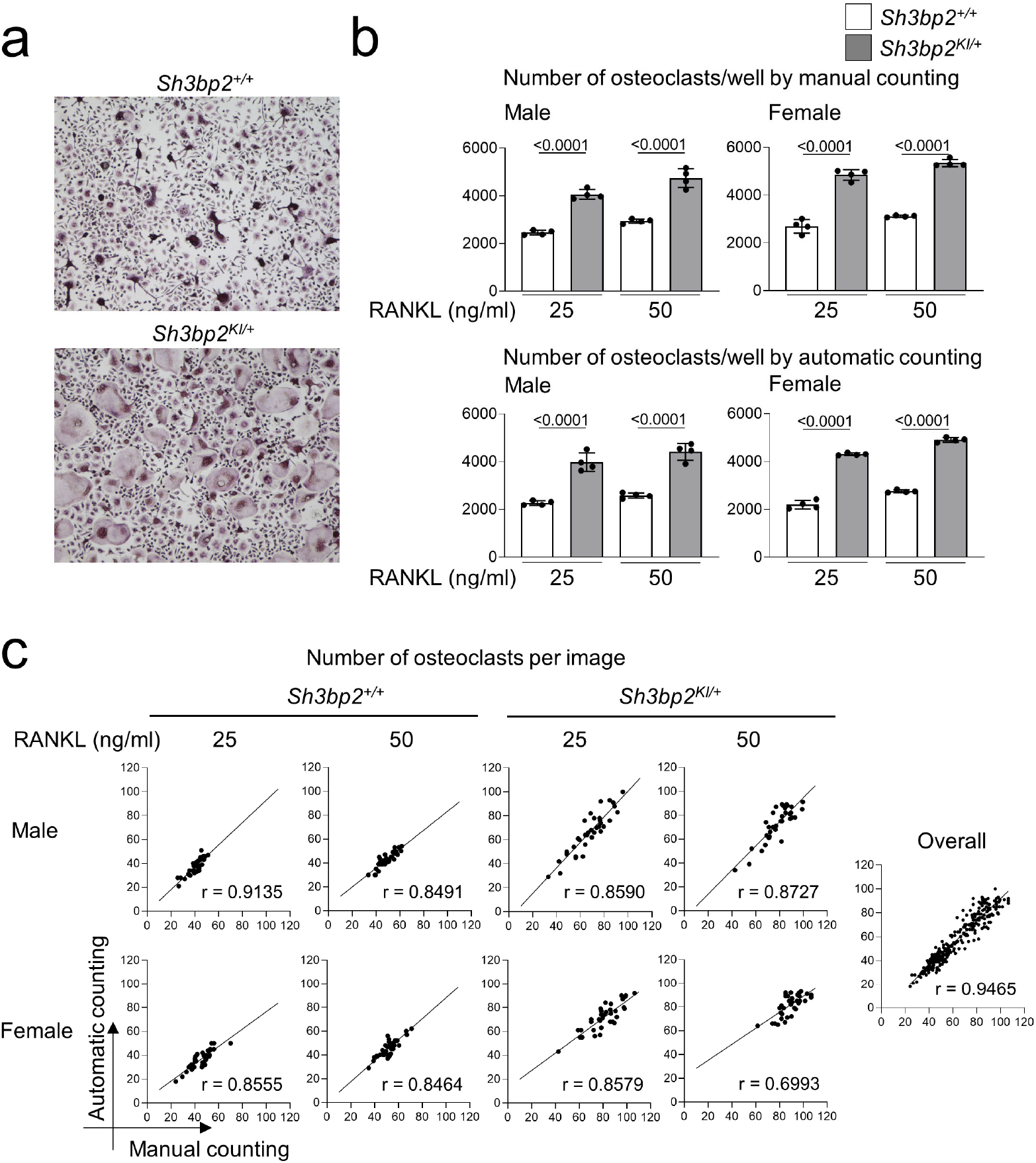
The automatic osteoclast counting compared with manual counting. **a,** Microscopic images of TRAP-stained osteoclast culture. Upper panel: *Sh3bp2^+/+^*; Lower panel: *Sh3bp2^KI/+^*. **b,** Validation of the performance of automatic osteoclast-counting in the practical situation. Upper panel: The number of osteoclasts measured manually as reference data. Lower panel: Number of osteoclasts measured by the automated system. Values on the graphs are *p* values calculated by the Tukey-Kramer test. The Y-axis is the number of osteoclasts per culture well, and the X-axis is concentration (25 and 50 ng/ml) of RANKL. **c** correlation analysis between automatic and manual osteoclast-counting. The number of osteoclasts on each image was analyzed manually and automatically. 36 images were analyzed for each condition. Pearson’s correlation coefficients (r) were shown, except for the analysis for overall samples, which did not show Gaussian distribution and was analyzed with Spearman’s analysis.

In Fig. 5c, we also compared the number of osteoclasts measured by a human examiner and OC_Finder for each of 36 microscopic images. In all the culture conditions, OC_Finder showed a high correlation, between 0.70 to 0.91, with manual counting (Fig. 5c left). The correlation was as high as r = 0.9465 when all samples were pooled and analyzed (Fig. 5c right). Thus, we confirmed that the automated counting system could generate comparable data to manual counting, and the system demonstrated a good sensitivity to detect biological differences in the experiment.

## Discussion

OC_Finder is the first fully automated osteoclast counting system that utilizes a deep-learning neural network. OC_Finder performs image segmentation and classification tasks in its pipeline. Overall, OC_Finder showed high accuracy in both tasks. When used for a practical scenario of counting osteoclasts (i.e. identifying and classifying cells) in microscopic images, OC_Finder showed comparable performance with human eyes (Fig. 5). Therefore, the system can provide valuable assistance in labor-intensive cell counting and greatly reduce the workload for researchers while maintaining acceptable recall and accuracy.

When the entire pipeline of OC_Finder was applied to microscopic images, the overall accuracy was affected by the segmentation step, which had a slightly lower accuracy than classification. Thus, improvement in segmentation will further increase the system’s accuracy, which is left as a future work. The quality of image segmentation may be controlled by changing parameters, such as the threshold, the filter size, and optimal values would be different for different input microscopic images. Users are encouraged to control the parameters for optimizing performance on their own dataset. The code for OC_Finder can be expanded for other similar cell classification tasks by re-training networks on a specific dataset. Expanding the method to handle other cell images is straightforward. OC_Finder will be able to extend to other similar works and be a widely used tool for cell image localization and detection.

## Acknowledgments

The authors are grateful to Jacob Verburgt for his help in preparing the manuscript. This work was partly supported by the National Institutes of General Medical Sciences (R01GM123055, R01GM133840) to DK, the National Science Foundation (CMMI1825941, MCB1925643, DBI2003635) to DK, and the National Institute of Dental and Craniofacial Research (R01DE025870, R01DE025870-06S1, R21DE030561) to YU.

## Author Contributions

MK conceived the study. MK cultured cells and collected the cell image data. XW developed the computational method, OC_Finder, and performed the computation. YH and YZ participated in the initial stage of the method development. XW, MK, YU, and DK analyzed the results. MK validated the performance of OC_Finder. MK and XW drafted the manuscript. DK edited the manuscript. YU participated in the editing process. DK administered the project. All authors have read and approved the manuscript.

## Corresponding author

Correspondence to Daisuke Kihara.

## Methods

### *In vitro* osteoclast differentiation

Bone marrow cells were isolated from the tibia, femur, and ilium of 7- to 8-week-old male and female *Sh3bp2^+/+^* and *Sh3bp2^KI/+^* mice on the C57BL/6 background. The *Sh3bp2^KI/+^* mice that have the heterozygous gain-of-function mutation in SH3-domain binding protein 2 (SH3BP2) were previously reported.^19,23,24^ After treating with red blood cell lysis buffer (eBioscience), bone marrow cells were cultured in alpha-MEM supplemented with 10% FBS and 1% penicillin/streptomycin on the Petri dish. After 3 hours, non-adherent cells were collected and further cultured in alpha-MEM containing 25 ng/ml M-CSF (PeproTech) on the Petri dish for three days to selectively grow the bone marrow-derived M-CSF-dependent macrophages (BMMs). BMMs were harvested, seeded on 48-well plates at the density of 2.5 × 10^4^ cells per well, and cultured for 3 days in the presence of four combinations of cytokines: 1) 25 ng/ml M-CSF and 25 ng/ml RANKL, 2) 25 ng/ml M-CSF and 50 ng/ml RANKL, 3) 25 ng/ml M-CSF and 50 ng/ml RANKL and 100 ng/ml TNF-α, 4) 25 ng/ml M-CSF and 50 ng/ml RANKL and 10 ng/ml IL-1β. To train the neural network with a variety of osteoclasts in size or morphology, two different BMMs sources (*Sh3bp2^+/+^* and *Sh3bp2^KI/+^* mice) in the presence of four different cytokine combinations were used. All cytokines were obtained from PeproTech.

#### Osteoclast and non-osteoclast image collection

Cells were stained by tartrate-resistant acid phosphatase (TRAP) staining (Sigma-Aldrich) and images were captured using the BZ-X810 microscopy (Keyence) with the following settings: 10x objective lens, 1/175 s exposure time, and 50 % transmitted light power. TRAP-positive cells containing more than 3 nuclei were regarded as osteoclasts. We obtained 458 microscopic images (314 images from *Sh3bp2^+/+^* cell culture and 144 images from *Sh3bp2^KI/+^* cell culture) that contain 13822 osteoclast and 13833 non-osteoclast images for the training, validation, and test of the neural network. The dataset included the same microscope images, i.e. 229 images each, from male and female. Osteoclasts and non-osteoclasts were identified and distinguished by visual evaluation of two independent individuals. The absolute coordinates of each osteoclast and non-osteoclast on the images were provided manually using the “Multi-point” function of ImageJ (NIH) and used to locate the osteoclasts and non-osteoclasts.

#### Counting the number of osteoclasts

After TRAP staining, nine images were captured (10X objective lens, 1/175 s exposure time, 50 % transmitted light power) from each well of the osteoclast culture. Osteoclasts in each of the nine images were identified and counted either by visual evaluation or by OC_Finder. To calculate the total number of osteoclasts per each culture well (for Fig. 5b), the numbers of osteoclasts per image of nine images from the well was averaged and normalized using the size of the area covered by a single image (1.587 mm^2^) and the surface area of the well (0.95 × 10^2^ mm^2^).

### Segmentation of cell images

Cell images are segmented from an input microscopic image using the watershed algorithm^7,8^. We chose this algorithm because it was successful in medical image segmentation tasks^26^. To improve the performance of the watershed algorithm, we applied the following pre-processing before applying watershed. Image were first converted to a grayscale image, then by binarized by Otsu’s method^6^, which roughly estimates the boundaries between foreground (cell regions) and background. Subsequently, we further applied morphological opening and closing operations to smoothen the cell regions. Otsu’s binarization determines a threshold value to distinguish foreground and background. This algorithm first computes the histogram of grayscales of pixels in an input image. Then, the algorithm applies different intensity thresholds to split the distribution to two distributions. For each threshold, it computes a weighted sum of variance of two distributions and the threshold that yielded the largest sum variance is selected to split foreground and background. This process is performed on-the-fly for each image, and thus no training process is needed.

Next, we removed noise in foreground regions by applying morphological opening and closing operations^22^ with a filter of a 3×3 pixel size. Opening operation in general removes irregular regions at the boundary and closing operation removes holes in foreground regions.

Subsequently, we applied a distance transform^27^ to all foreground regions, which tries to separate individual cells from large foreground regions that include multiple cells. In the distance transform, the label of each pixel, which is binary, 0 or 1, at this step due to the Otsu’s binarization, was updated to the distance to the closest background pixel. Thus, pixels that are deep inside a cell tend to have a large value. We used 0.7 of the maximum distance in the entire image as the threshold to select pixels as possible cell centers (called markers), which were used as starting points by the watershed algorithm. This procedure was implemented with the OpenCV^28^ package.

### Data Augmentation for network training

During training the network, we augmented input cell images by randomly applying one of 12 image transformations^29,30^. A magnitude of a transformation was also randomly chosen from a pre-defined range. We followed AutoAugment^30^ to decide the types and the magnitude range of transformations to apply. The 12 types of transformations and the magnitude range are listed in Supplementary Table 1. Examples of the 12 augmentation types are shown in Supplementary Fig. 3.

### Training the deep neural network

We used ResNet18^21^ in this work. We also tried to use deeper ResNet networks but did not observe clear improvement (data not shown). Out of 458 microscopic images, 81.4% (373) were used for training and the rest of 18.6% (85) of them were used for testing. The training set was further split into two parts, 80% (298) used for training and 20% (75) for validation. Thus, the data split was performed with microscopic images but the classification was performed at the individual extracted cell image level. The number of non-osteoclasts and osteoclasts included in the training, validation, and testing are 9276/9278, 2219/2226, and 2327/2329 respectively for non-osteoclast/osteoclasts. These cells were manually labeled and the numbers do not include small cell regions with less than 500 pixels.

RGB values of a pixel in an image in the training set were normalized by computing the Z-score:

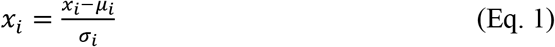

where *x_i_* is the *i*th channel (R, G, or B) of a pixel *x_i_ μ_i_* and *σ_i_* are the mean and the variance of channel *i* in the training set, *σ_i_*. In the validation and testing stages, we used the same mean and variance values that were taken from the training set. In training, each input cell image was subject to an augmentation. The type and the magnitude of augmentation were randomly selected.

We used a teacher-student architecture, Mean-Teacher^16^ for training the model because it is in general effective in avoiding overfitting. The Mean-Teacher model^16^ updates weights of a teacher model with a moving average of the weights from a sequence of student models as

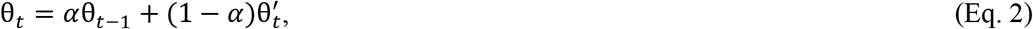

where 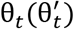 is the parameters of the Teacher (Student) Model at update step t. *α* is a smoothing coefficient. We tried different *α* as shown in Supplementary Table 3 and set it to 0.999 as it gave the highest accuracy in the validation. We used the teacher model in the evaluation.

Two parameters, regularization parameters of L2 regularization we used and the learning rate were optimized with the Adam Optimizer^31^ for minimizing a cross entropy loss. The regularization parameter values tested were [1e-7, 1e-6, 1e-5, 1e-4, 1e-3, 1e-2, 1e-1] and the learning rate values tested were [2e-5, 2e-4, 2e-3, 2e-2, 2e-1, 2]. Based on the performance on the validation set, a regularization parameter of 1e-5 and a learning rate of 0.002 performed the best. Under a hyperparameter combination, we generated 100 trained models trained on the training set, which were kept at each epoch. Among them, we selected the model with the above-mentioned best hyperparameter combination, which performed the best on the validation set and applied it to the test set. The batch size was set to 256 images and the models were trained for 500 epochs.

### Statistical analysis

One-way ANOVA with Tukey-Kramer post hoc test was used for the comparison between groups. For correlation analysis, Pearson’s correlation analysis was used for the samples with Gaussian distribution. For the samples that did not show Gaussian distribution, Spearman’s analysis was used. GraphPad Prism (ver.7; GraphPad Software, La Jolla, CA) was used for all statistical analyses.

### Reporting summary

Further information on research design is available in the Nature Research Reporting Summary linked to this article.

### Data availability

The labeled data for training and verifying our methods can be downloaded from https://doi.org/10.5281/zenodo.5022015. More data that support the findings of this study are available from the corresponding author upon request.

### Code availability

The OC_Finder program is freely available for academic users at http://github.com/kiharalab/OC_Finder.

## Supplementary Information

**Supplementary Table 1.** Transformation details for training

| **Transformation Name** | **Description** | **Range** |
| --- | --- | --- |
| TranslateX(Y) | Translate the image in the horizontal (vertical) direction by magnitude number of pixels. | [-0.3,0.3] |
| Rotate | Rotate the image magnitude degrees. | [-30,30] |
| AutoContrast | Maximize the image contrast, by making the darkest pixel black and lightest pixel white. | [0 or 1] |
| Invert | Invert the pixels of the image | [0 or 1] |
| Equalize | Equalize the image histogram | [0 or 1] |
| Solarize | Invert all pixels above a threshold value of magnitude. | [0, 256] |
| Posterize | Reduce the number of bits for each pixel to magnitude bits | [4,8] |
| Contrast | Control the contrast of the image. A magnitude=0 gives a gray image, whereas magnitude=1 gives the original image. | [0.1,1.9] |
| Color | Adjust the color balance of the image, in a manner similar to the controls on a color TV set. A magnitude=0 gives a black & white image, whereas magnitude=1 gives the original image. | [0.1,1.9] |
| Brightness | Adjust the brightness of the image. A magnitude=0 gives a black image, whereas magnitude=1 gives the original image. | [0.1,1.9] |
| Sharpeness | Adjust the sharpness of the image. A magnitude=0 gives a blurred image, whereas magnitude=1 gives the original image | [0.1,1.9] |

**Supplementary Fig. 1.**
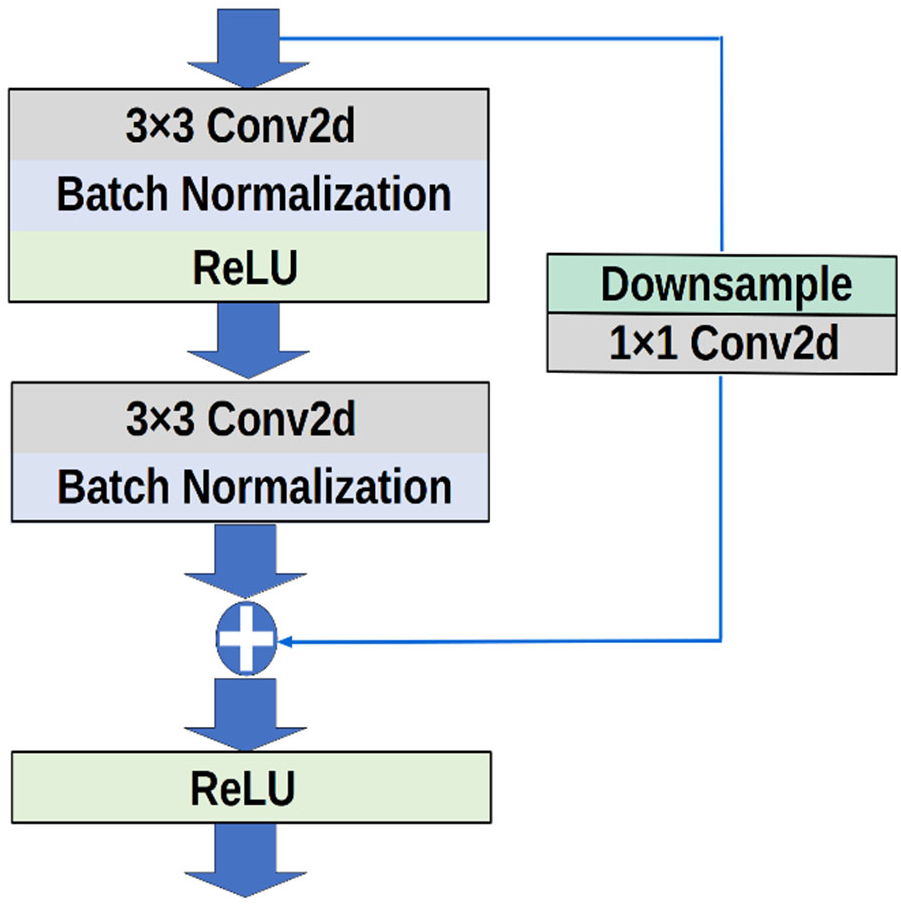
The architecture of the residual block (ResBlock). For a given input image patch, a convolutional filer with size of 3*3, batch normalization, and a ReLU activation are sequentially applied. To further aggregate the spatial information, another convolutional filter and batch normalization are applied for the first output. To avoid the information loss of initial input, a downsample module with a convolutional filter with a size of 1×1 is applied to the initial input to reduce the size of feature map. The network we used has 4 residual layers with 64, 128, 256, and 512 residual blocks, respectively (Fig. 1b). Among them, down-sampling is adopted for the last 3 residual layers. If no down-sampling is applied, an identity mapping will apply to the input to output. These two outputs, residual output and the first input, were added and passed to a ReLU activation.

**Supplementary Table 2.** Cell segmentation results on the 10 microscopic images.

| **Image** | **Total** | **Not Identified** | **Detection Rate (%)** | **Additional Detections by OC-Finder** |
| --- | --- | --- | --- | --- |
| Female *Sh3bp2*<sup>+/+</sup> stimulated with RANKL (25 ng/ml) | 1723 | 22 | 98.7 | 0 |
| Female *Sh3bp2*<sup>+/+</sup> stimulated with RANKL (50 ng/ml) | 1823 | 17 | 99.1 | 0 |
| Female *Sh3bp2*<sup>KI/+</sup> stimulated with RANKL (25 ng/ml) | 1014 | 5 | 99.5 | 0 |
| Female *Sh3bp2*<sup>KI/+</sup> stimulated with RANKL (50 ng/ml) | 883 | 6 | 99.3 | 0 |
| Male *Sh3bp2*<sup>+/+</sup> stimulated with RANKL (25 ng/ml) | 1053 | 18 | 98.3 | 0 |
| Male *Sh3bp2*<sup>+/+</sup> stimulated with RANKL (50 ng/ml) | 1204 | 7 | 99.4 | 0 |
| Male *Sh3bp2*<sup>KI/+</sup> stimulated with RANKL (25 ng/ml) | 445 | 0 | 100.0 | 3 |
| Male *Sh3bp2*<sup>KI/+</sup> stimulated with RANKL (50 ng/ml) | 664 | 2 | 99.7 | 0 |
| Male *Sh3bp2*<sup>+/+</sup> stimulated with RANKL (50 ng/ml) and TNF-α | 635 | 3 | 99.5 | 0 |
| Male *Sh3bp2*<sup>+/+</sup> stimulated with RANKL (50 ng/ml) and IL-1β | 777 | 0 | 100.0 | 0 |
| **Average** | 1022.1 | 8 | 99.4 | 0.3 |

Cell segmentation by OC\_Finder was compared with manually identified cells in 10 microscopic images. The image column lists specification of each images. Total, the total number of cells that were manually identified in each image data. Not identified, the number of cells that were not detected by OC\_Finder. Detection rate was computed by (Total – Not\_Identified)/Total \* 100(%). Additional detection by OC\_Finder, the number of additional segments OC\_Finder identified that did not correspond to manual identification.

**Supplementary Table 3.**
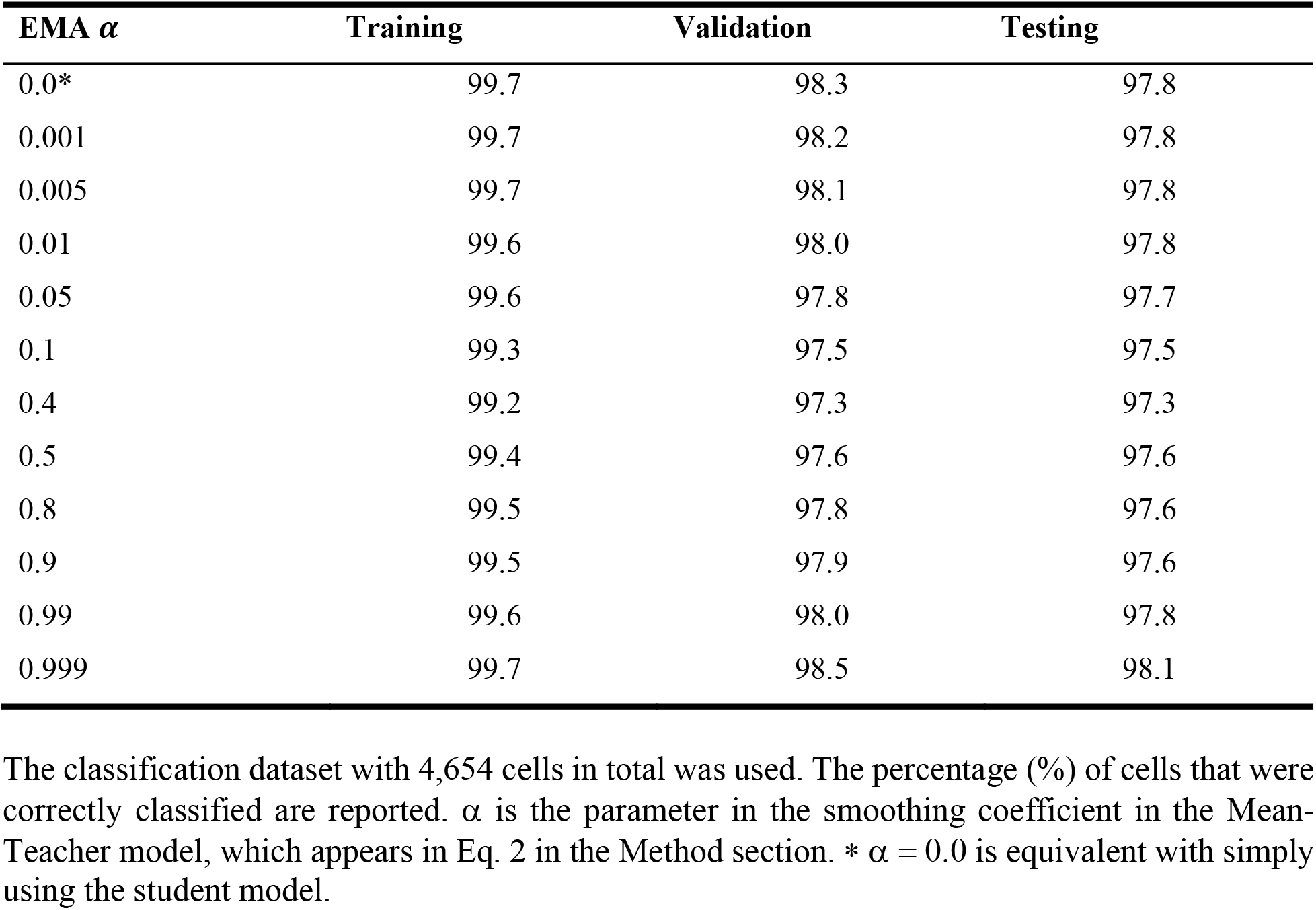
Accuracy of the Mean-Teacher model with different smoothing coefficient values (α).

| EMA $\alpha$ | Training | Validation | Testing |
| --- | --- | --- | --- |
| 0.0* | 99.7 | 98.3 | 97.8 |
| 0.001 | 99.7 | 98.2 | 97.8 |
| 0.005 | 99.7 | 98.1 | 97.8 |
| 0.01 | 99.6 | 98.0 | 97.8 |
| 0.05 | 99.6 | 97.8 | 97.7 |
| 0.1 | 99.3 | 97.5 | 97.5 |
| 0.4 | 99.2 | 97.3 | 97.3 |
| 0.5 | 99.4 | 97.6 | 97.6 |
| 0.8 | 99.5 | 97.8 | 97.6 |
| 0.9 | 99.5 | 97.9 | 97.6 |
| 0.99 | 99.6 | 98.0 | 97.8 |
| 0.999 | 99.7 | 98.5 | 98.1 |

The classification dataset with 4,654 cells in total was used. The percentage (%) of cells that were correctly classified are reported. $\alpha$ is the parameter in the smoothing coefficient in the Mean-Teacher model, which appears in Eq. 2 in the Method section. \* $\alpha = 0.0$ is equivalent with simply using the student model.

**Supplementary Fig. 2.**
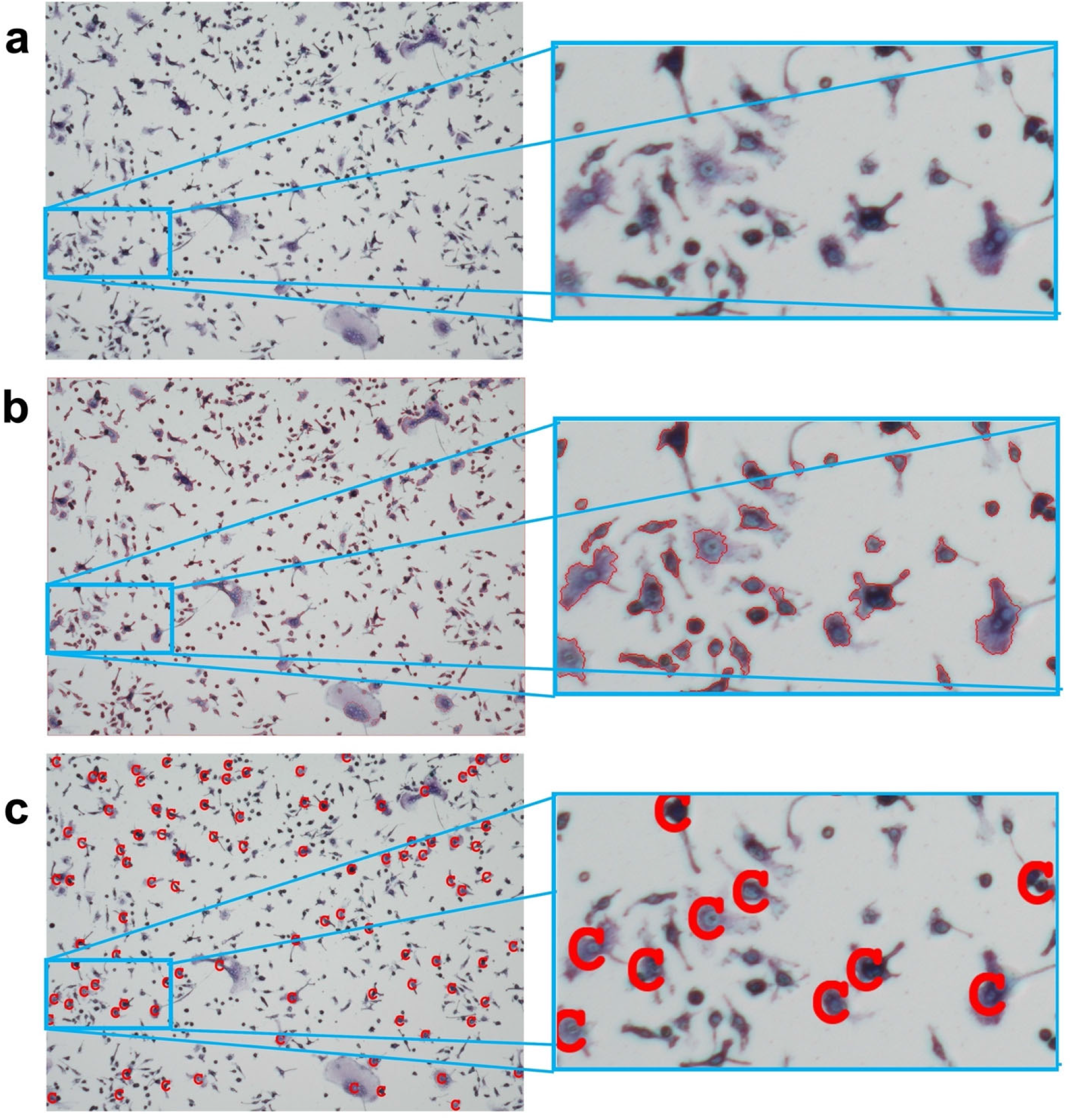
Example of segmentation. **a.** The input image from BZ-X810 microscopy. **b.** Segmented areas by the Watershed algorithm. The boundary is shown in red. **c.** Center locations (shown with C) of the filtered segmented areas, which are cropped with 50×50 pixel size squares. These cropped squares are input images for the deep learning model for cell classification.

**Supplementary Fig. 3.**
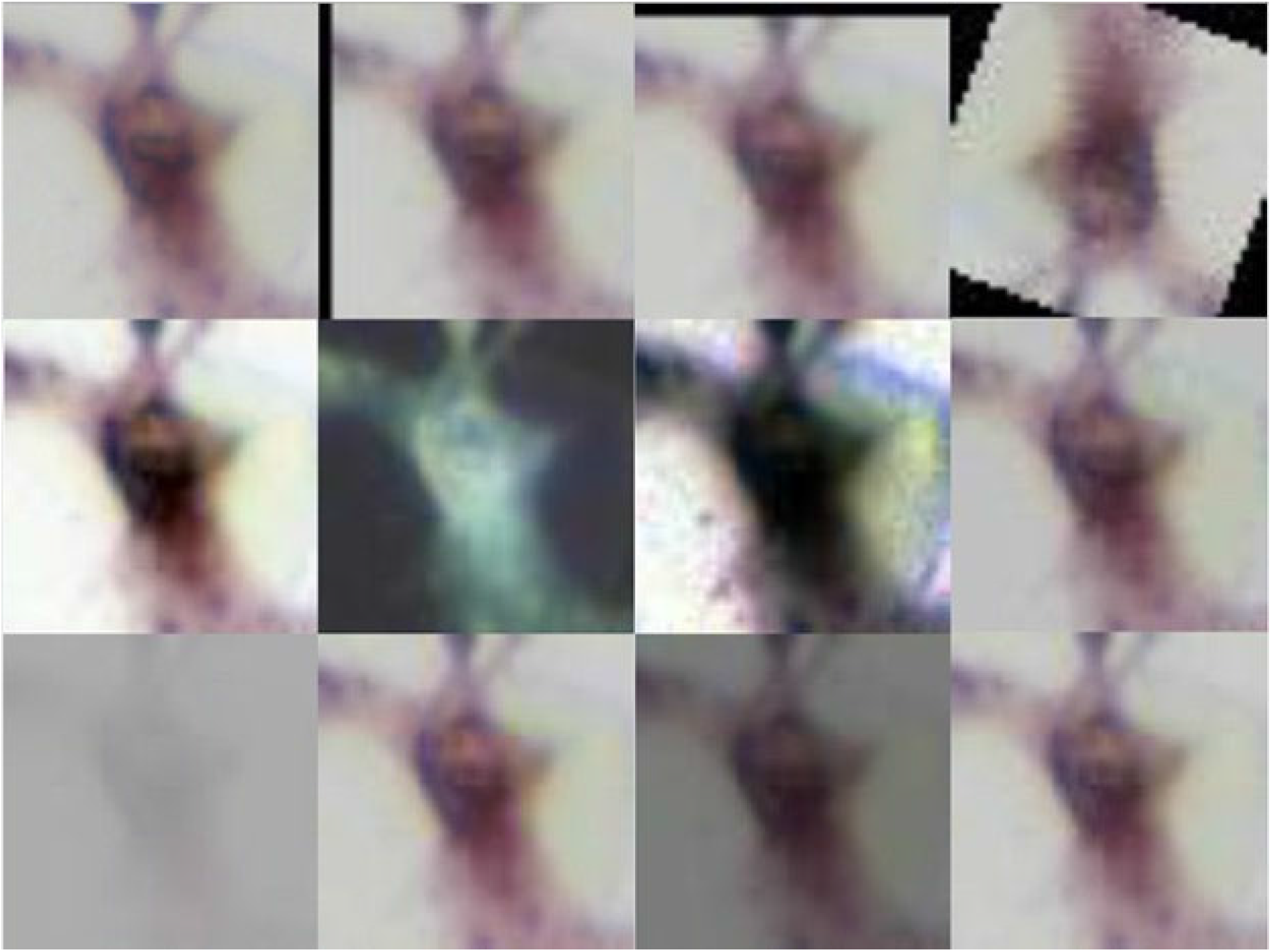
Examples of 12 different augmentations. From the top to bottom rows, left to right: the original image, image with translate-X, translate-Y, rotate, auto-contrast, invert, equalize, solarize, posturize, contrast, color, brightness, and sharpness.

